# Allometric Scaling of physiologically-relevant organoids

**DOI:** 10.1101/559682

**Authors:** Chiara Magliaro, Andrea Rinaldo, Arti Ahluwalia

**Affiliations:** Research Center “E. Piaggio”, University of Pisa, Largo Lucio Lazzarino, 1, 56122, Pisa, Italy; Laboratory of Ecohydrology, Ecole Polytechnique Fédérale de Lausanne, Lausanne 1015, Switzerland and Dipartimento ICEA, University of Padova, via Loredan 30, 35131 Padova, Italy; Department of Information Engineering, University of Pisa, Via Caruso, 16, 56122, Pisa, Italy

**Keywords:** *in silico* modelling, brain organoids, liver organoids, oxygen consumption, allometric scaling, physiological relevance, micro-fluidics

## Abstract

The functional and structural resemblance of organoids to mammalian organs suggests that they might follow the same allometric scaling rules. However, despite their remarkable likeness to downscaled organs, non-luminal organoids are often reported to possess necrotic cores due to oxygen diffusion limits. To assess their potential as physiologically relevant *in vitro* models, we determined the range of organoid masses in which quarter power scaling as well as a minimum threshold oxygen concentration is maintained. Using data on brain organoids as a reference, computational models were developed to estimate oxygen consumption and diffusion at different stages of growth. The results show that mature brain (or other non-luminal) organoids generated using current protocols must lie within a narrow range of masses to maintain both quarter power scaling and viable cores. However, micro-fluidic oxygen delivery methods could be designed to widen this range, ensuring a minimum viable oxygen threshold throughout the constructs and mass dependent metabolic scaling. The results provide new insights into the significance of the allometric exponent in systems without a resource-supplying network and may be used to guide the design of more predictive and physiologically relevant *in vitro* models, providing an effective alternative to animals in research.

---

Recent advances in *in vitro* technology have led to the development of organoids, i.e. cell aggregates grown from a small number of stem cells able to self-assemble, recapitulating the three-dimensional architecture of an organ at the micro-scale^1,2^. They are currently considered as one of the most promising ways to study cell behaviour and may have significant potential as a tool for investigating developmental biology, disease pathology, regenerative medicine and drug toxicity. Moreover, their downscaled functional and structural resemblance to mammalian organs suggests that they might also follow allometric scaling rules, such as Kleiber’s law (KL)^3–5^.

KL is one of the most well-known allometric relationships in biology. It states that the basal metabolic rate (*BMR*) of an organism scales with its body mass, *M*, according to a universal power-law *BMR* = *aM^α^*, (where *α* is a scaling exponent, *a* a constant). On the basis that the number of cells in an organism is proportional to its mass, an analogous expression relates the average resource consumption rate per cell (cellular metabolic rate, *CMR*) to mass as *CMR ∝ M*^(*α−1*)^ =*M^b^*. The majority of experimental studies support the claim that KL applies across species with *α*=3/4, or *b*=−1/4 hence it is often referred to as the quarter power scaling law^6^. A number of theoretical explanations have been proposed to support quarter power scaling^5,7^. As KL applies across several orders of magnitude of mass, it has been described as a unifying equation of biology^8^. Indeed, metabolic requirements of organisms, as well as their constituent cells, influence many fundamental biological properties common to all forms of life such as growth, differentiation and death. The pertinence of KL to the development of physiologically relevant *in vitro* models has been highlighted^9,10^, and it is considered by some scientists as the key to enabling the extrapolation of biological parameters from micro-scaled in *vitro* systems to the *in vivo* context: one of the holy grails of modern biomedical science^11^.

Ahluwalia^12^ recently demonstrated that cells *in vitro* can maintain KL even the in absence of a resource-supplying network. In particular, given that oxygen is the primary metabolic resource for cells, the author evaluated the *CMR* of cell spheroids with increasing radii using computational mass transfer models coupling Michaelis-Menten (MM) reaction kinetics and oxygen diffusion through 3D constructs. The results show that KL holds in 3D spheroids if they have a suitable mass and cell density to enable the formation of a sufficiently large concentration gradient within the construct. However, the model proposed by Ahluwalia did not focus on organoid technology as it did not take into account the fact that organoids generated from embryoid bodies rely on the use of a gelatinous protein mixture resembling the extracellular environment surrounding the construct (e.g., Matrigel or Geltrex). While the 3D extra-cellular matrix (ECM) mimic supports cell survival and function^13–15^, it might be a limiting factor for oxygen uptake within the structure, particularly in the case of non-luminal *in vitro* replicates such as brain, liver or pancreatic organoids. Moreover, the cell density of standard *in vitro* cultures considered in the paper (i.e. 5.14·10^12^ cells/m^3^) is about two orders of magnitude lower than those typically reported for such organoids (around 10^14^ cells/m^3^) which are close to physiological cell densities^1,13,14,16^. Thus, for a fixed mass or radius, the oxygen consumption rate per unit volume of non-luminal organoids is much higher than in spheroids, which often leads to a dead core - a recurring feature in these systems^17^. By combining computational models with images of mature brain organoid slices stained with a nuclear dye, Berger *et al*.^17^ recently identified a minimum threshold oxygen concentration for viability, *C_crit_*, of 0.04 mol/m^3^, which is also the critical value needed for mitochondrial ATP production^18^.

The scientific community is aware of the urgent need to improve organoid technology in terms of quality and reproducibility of sample size and shape. Particular attention should be addressed to eliminate, or at least minimise, the ‘non-viable’ centres while maintaining the ability of stem cells to self-organize and differentiate recapitulating physiological forms and functions^19–22^. This study is thus aimed at identifying a working window in which two criteria necessary for physiological relevance in cell-filled (non-luminal) organoids are met: i) they follow “physiological” quarter power scaling; and ii) the oxygen concentration within their volume is maintained above the critical threshold *C_crit_*. By customising the computational framework based on allometric scaling outlined by Ahluwalia^12^, the *CMR* of spherical organoid models was computed over 5 orders of magnitude of mass to estimate the value of the exponent *b*. Additionally, the concentration gradient through the models was mapped to determine their minimum oxygen levels. Finally, to investigate the effects of integrating vasculature in organoids, a single fluidic channel was added to the models.

## Computational methods

To compute the oxygen concentration in organoid systems and determine the allometric relationships between the cellular metabolic rate (CMR) and organoid mass, 3D multi-physics models were implemented and solved numerically using COMSOL Multiphysics (version 4.3 COMSOL AB, Stockholm, Sweden) and the UMFPACK direct solver. In the models, reaction, convection and diffusion of oxygen through organoids and gels as well as fluid dynamics for incompressible fluids were considered.

### Organoid volume expansion and stem cell proliferation

Referring to published data on brain organoids derived from human induced pluripotent stem cells (hiPSCs)^13,17^, we modelled both stem cell proliferation and organoid volume expansion to identify a range of experimental organoid masses and corresponding cell densities. In particular, cell proliferation was modelled using the Sherley model^23^:

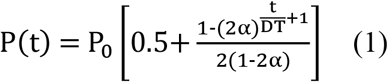

where P(t) is the population size at time t, P_0_ the initial number of cells, α the mitotic fraction and DT is the division time. Since we supposed that all the cells within the organoid reach mitosis, α was set to 1. The division time, DT = 1.19·10^5^ s, was estimated from the data reported in Berger *et al*.^17^, where 9000 hiPSCs were plated and a total of 1.8·10^5^ cells were counted at the end of the experiment after 20 days.

On the other hand, the function describing organoid expansion was extracted by least squares fitting the data on measurements of organoid diameter reported Monzel *et al*.^13^ to a generic exponential:

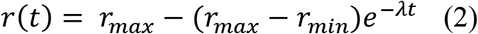

where r(t) is the organoid radius at time t, r_min_ the initial radius (i.e. 250 μm) and r_max_ the maximum radius reached at the end of its growth (i.e. 562 μm). The expansion coefficient, λ, was estimated as 1.126·10^−5^ s^−1^ (R^2^: 0.9884, RMSE: 16.43).

From the equations above, we calculated the cell densities at three time points (i.e. day 4, day 8 and day 12, Table 1). Since the cell densities vary by less than 20% over time, a mean value of 2.52·10^14^ cells/m^3^ was used for all three time points. This value is of the same order of magnitude as those reported in other protocols for non-luminal organoid generation (e.g., ref.^24^).

**Table 1.** Number of cells, average organoid radii, mass and cell densities at each time point investigated. Data on hiPSC generated brain organoids from Monzel *et al*.^13^ and Berger *et al*.^17^.

| Time [day] | Number of cells | Radius [$\mu\text{m}$] | Mass [mg]* | Cell density [cells/$\text{m}^3$] |
| --- | --- | --- | --- | --- |
| 4 | 9000 | 250 | 0.0654 | $1.35 \cdot 10^{14}$ |
| 8 | 66300 | 399 | 0.2666 | $2.47 \cdot 10^{14}$ |
| 12 | 180000 | 562 | 0.7439 | $3.75 \cdot 10^{14}$ |

\*Organoid density = $1000 \text{ kg}/\text{m}^3$

When developing the computational models, four basic phenomena were considered: i) cell proliferation, ii) organoid volume expansion, iii) oxygen consumption within the organoid and iv) oxygen diffusion. Given that the characteristic times for organoid growth and expansion are much longer than the times for oxygen diffusion and consumption within the spheroid (Table 2), we modeled each radius and time point (4, 8 and 12 days respectively) as a quasi-steady-state model of reaction and diffusion.

**Table 2.** Characteristic times for the phenomena considered.

| Phenomenon | Equation | Characteristic time [s] |
| --- | --- | --- |
| Oxygen Diffusion within the organoid | $\frac{r_{max}^2}{D}$ | $\sim 336$ |
| Oxygen consumption | $\frac{k_m + c_0}{OCR \rho}$ | $\sim 50$ |
| Organoid volume expansion | $\frac{1}{\lambda}$ | $\sim 8.8 \cdot 10^4$ |
| Stem cell proliferation | $DT$ | $1.19 \cdot 10^5$ |

### The model geometries

The model geometries are divided into three sub-domains: i) the organoid, represented by a solid sphere where both oxygen diffusion and consumption are solved, ii) a shell of hydrogel surrounding the organoid domain, whose thickness varies maintaining a constant shell volume as the organoid grows in size, and where only oxygen diffusion is solved and iii) a cylinder with a diameter of 20 μm representing a central micro-channel through which medium flows and where both oxygen diffusion and fluid dynamics are solved. The sub-domains were combined stepwise to implement three different configurations (Figure 1) representing: (A) a cell-filled organoid model similar to the system reported in Ahluwalia^12^; (B) an organoid as in (A), surrounded by a shell of hydrogel; (C) a “vascularized” organoid as in (A) surrounded by a hydrogel shell.

**Figure 1.**
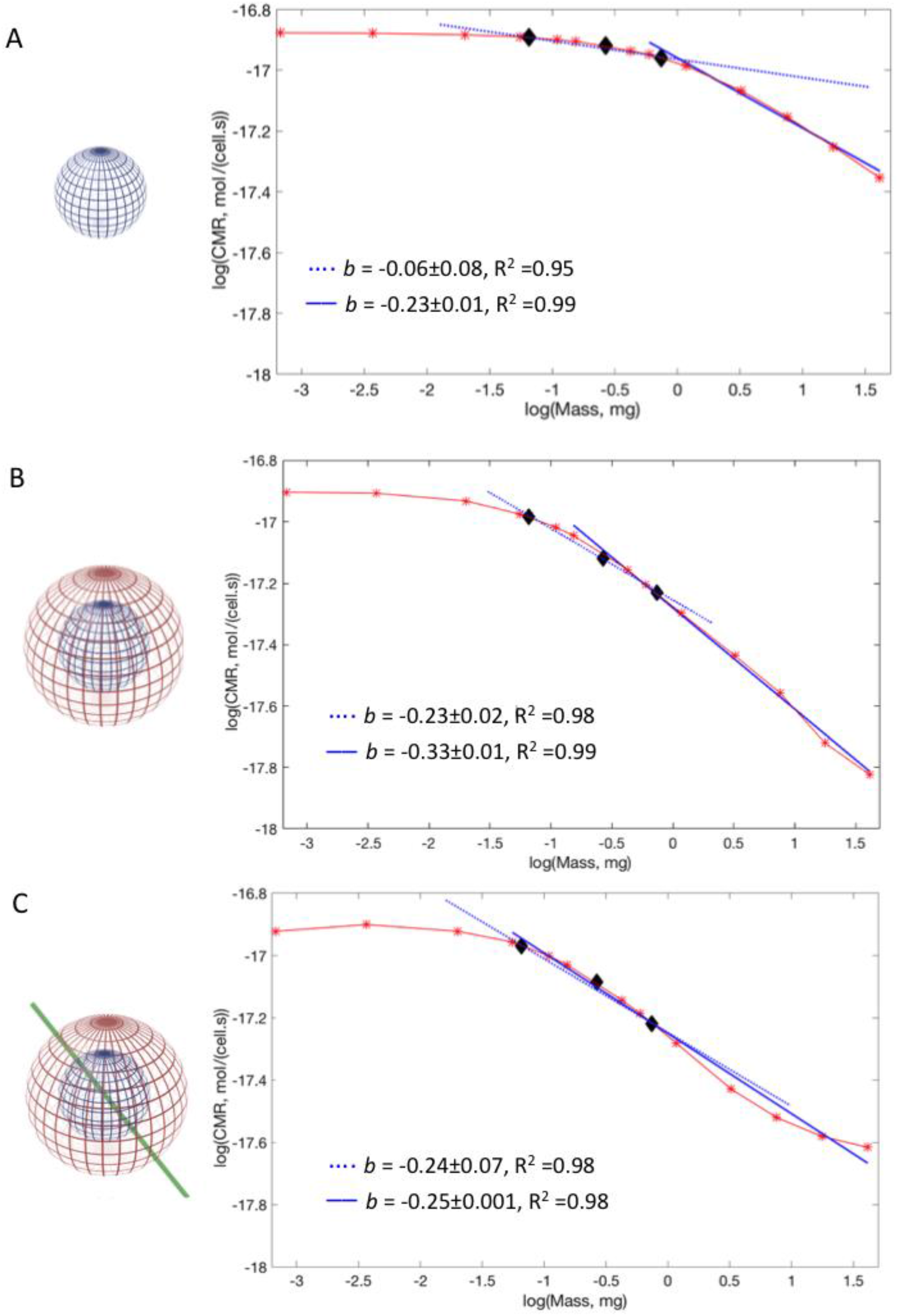
Model representations and computed *CMR* (red asterisks (*)) against mass for A) shell-free constructs; B) constructs with a hydrogel shell; C) constructs with a hydrogel shell and a micro-vessel. The experimental range of organoid masses in Table 1 are black diamonds (♦). Blue dotted line: least squares fit for the experimental range of masses. Blue full line: best fit. The slopes (*b*) and corresponding R^2^ values are indicated in the plots.

Cerebral organoids are not perfect spheres, but slightly ellipsoidal. In Figures 1A and B, the spherical constructs can be considered as the worst case in terms of surface area to volume ratio - and hence minimum oxygen concentration. On the other hand, the model in Figure 1C represents a structure with a blood vessel mimic, similar to what might occur in implanted cortical organoids after they connect to a host vasculature^21^.

### The multi-physics models

A reaction and diffusion model^25^ was implemented for mapping oxygen in the basic configuration shown in Figure 1A. Since organoid cell densities (*ρ*) are close to physiological (5.14·10^14^ cell/m^3^ *in vivo*^26^, versus 2.52·10^14^ cells/m^3^ from Table 1), we assumed that the oxygen diffusion constant within the organoid (D_org_) is the same as that in *in vivo* tissues^27^. Oxygen consumption was described using MM kinetics, considering literature values of *k_m_* and maximal oxygen consumption rate (OCR) typical for human stem cells (see Table 3).

**Table 3.**
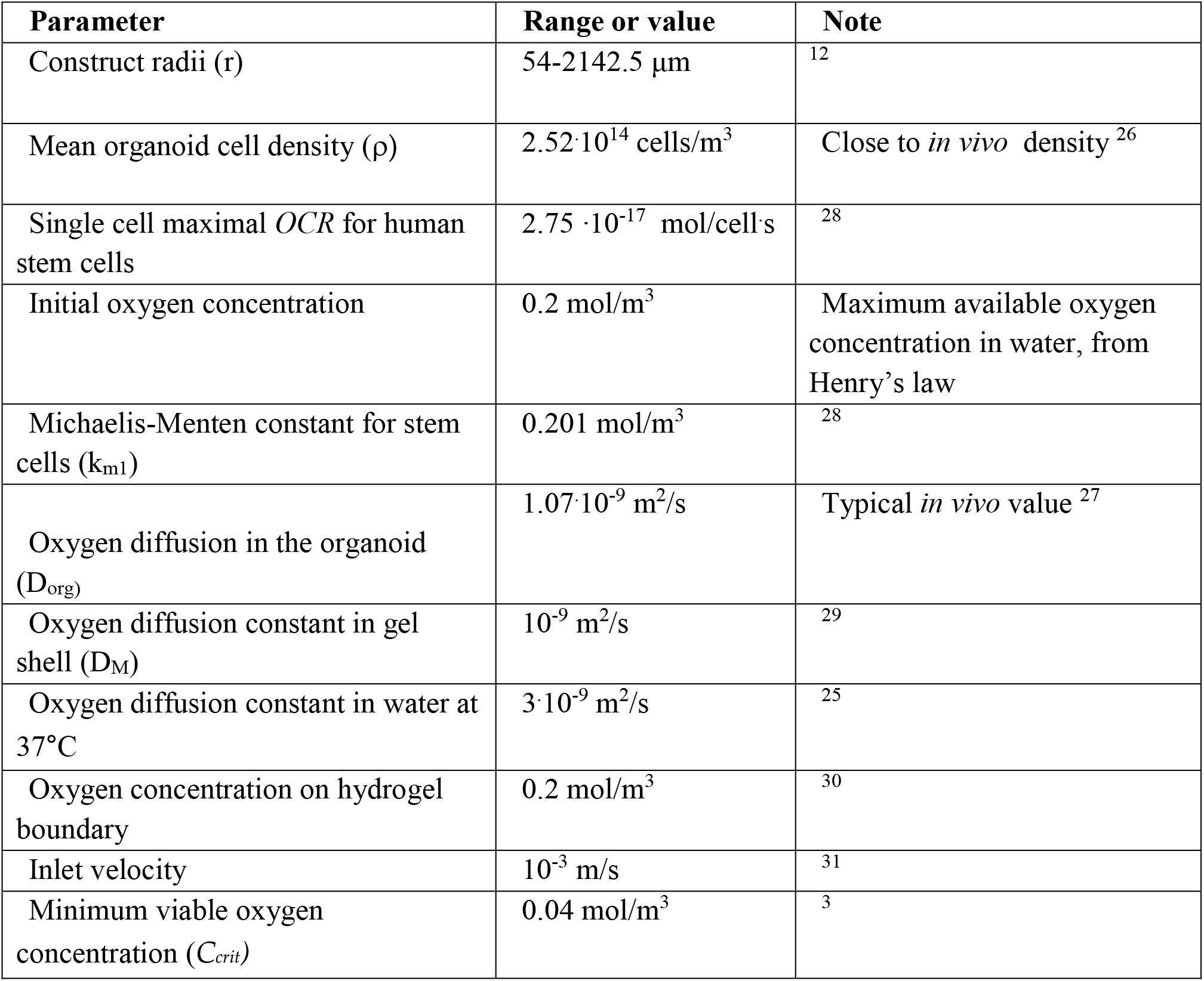
Parameter inputs to the computational models.

| Parameter | Range or value | Note |
| --- | --- | --- |
| Construct radii (r) | 54-2142.5 $\mu\text{m}$ | <sup>12</sup> |
| Mean organoid cell density ($\rho$) | $2.52 \cdot 10^{14}$ cells/$\text{m}^3$ | Close to *in vivo* density <sup>26</sup> |
| Single cell maximal *OCR* for human stem cells | $2.75 \cdot 10^{-17}$ mol/cell·s | <sup>28</sup> |
| Initial oxygen concentration | 0.2 mol/$\text{m}^3$ | Maximum available oxygen concentration in water, from Henry's law |
| Michaelis-Menten constant for stem cells ($k_{m1}$) | 0.201 mol/$\text{m}^3$ | <sup>28</sup> |
| Oxygen diffusion in the organoid ($D_{org}$) | $1.07 \cdot 10^{-9}$ $\text{m}^2/\text{s}$ | Typical *in vivo* value <sup>27</sup> |
| Oxygen diffusion constant in gel shell ($D_M$) | $10^{-9}$ $\text{m}^2/\text{s}$ | <sup>29</sup> |
| Oxygen diffusion constant in water at 37°C | $3 \cdot 10^{-9}$ $\text{m}^2/\text{s}$ | <sup>25</sup> |
| Oxygen concentration on hydrogel boundary | 0.2 mol/$\text{m}^3$ | <sup>30</sup> |
| Inlet velocity | $10^{-3}$ m/s | <sup>31</sup> |
| Minimum viable oxygen concentration ($C_{crit}$) | 0.04 mol/$\text{m}^3$ | <sup>3</sup> |

The boundary oxygen concentration was fixed at 0.2 mol/m^3^, assuming a well-mixed supply of medium to the organoids. The ruling equation for this system is:

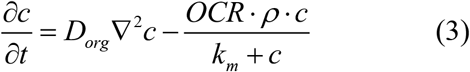

More detailed equations and boundary conditions are reported in the Supplementary Materials. In the second configuration in Figure 1B, only diffusion was implemented within the hydrogel shell. The oxygen diffusion constant in the shell, D_M_, was equal to that measured for Matrigel^29^. The cell-filled core was identical to the basic model in Figure 1A.

Finally, in the third configuration, oxygen mass transport considering diffusion, convection and consumption in the spheres (see Supplementary Materials) were coupled to fluid dynamics in the central channel based on the geometry shown in Figure 1C. The velocity field in the channel was solved using the Navier-Stokes equation for Newtonian fluids, setting the inlet velocity to 10^−3^ m/s, a value consistent with blood velocity within capillaries^31^. The simulations were run on constructs with the same radii as in ref.^12^ (ranging from 54.5 to 2142 μm) and includes the experimental radii reported in Table 1. All the constants and variable ranges used are listed in Table 3.

### Data evaluation and fitting

Once the solutions were obtained, the minimum oxygen concentration within the organoid domain was computed. In addition, the surface integration function on the organoid domain was used to determine the total inward oxygen flux at the organoid boundaries and the resulting *CMR* as in Ahluwalia^12^ Specifically, the average metabolic rate per cell or *CMR* is given by the surface integral of the inward oxygen flux, divided by the number of cells in the construct.

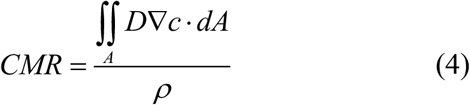

In the case of the microfluidic model, the positive outward oxygen flux from the walls of the central channel was subtracted from the negative inward flux at the surface of the organoid domain (further equations and details on the calculation of *CMR* are reported in the Supplementary Materials).

Both the *CMR* data and the minimum oxygen concentrations were then imported into Matlab 2017a (The Mathworks Inc., Boston Massachusetts) for plotting and curve fitting. To investigate the allometric behaviour of the constructs, log(CMR) versus log(mass) graphs were generated for each configuration using the method described in ref.^12^ and outlined in the Supplementary Materials. The allometric exponent b was estimated from the slope of the log-log curves for the range of experimental masses in Table 1. The best fit value of *b*, corresponding to the slope covering the maximum number of points and with the maximum R^2^ value, was also determined. Here, the log(*CMR*) - log(mass) dataset for each configuration was fitted to a straight line discarding one point at a time beginning from the lowest mass. A Gaussian equation was used to fit the minimum oxygen concentration data for each configuration implemented. The equations were used to estimate the radius at which the oxygen concentration is equal to *C_crit_* in the organoid domain.

## Results

As shown in Figure 1A, for gel-free models with small masses, the *CMR* is mass-independent (i.e. the slope *b*=0). This implies that all the cells in the constructs consume metabolic resources at the same rate since they all perceive the same oxygen concentration^12^. In the range of experimental organoid masses reported in Table 1, the slope *b*= −0.06 (R^2^: 0.95), which is far from quarter power scaling. Larger masses (corresponding to radii between 600 and 2000 μm) have a slope close to the KL value, *b*= −0.23 (R^2^: 0.99, best fit). The model suggests that a resource-supplying network is not a necessary condition for quarter power scaling, but allometric scaling will not occur below a minimum construct mass.

The constructs surrounded by a shell also have a constant *CMR* for small masses, although its value is slightly lower than that of the gel-free ones (Figure 1B versus 1A). This is reasonable because the oxygen concentration at the surface of the organoid domain with a gel shell is always less than 0.2 mol/m^3 12^. In the experimental range of interest *b* is close to −1/4 as in KL (*b*= −0.23, R^2^: 0.98), falling to *b*=−0.33 (R^2^: 0.99, best fit) for larger spheres. In the case of the constructs with the hydrogel shell and the micro-channel, the log-log graph for *CMR* versus mass (Figure 1C) shows that the experimental range of masses also follow KL as the slope *b* is almost −1/4. Moreover, *b* is still mantained within the allometric range for larger masses (*b*=−0.25, R^2^: 0.98).

Figure 2 reports the minimum oxygen concentration values, *C_min_*, within the organoid domains as a function of their radius and the fitted Gaussian curves for the three configurations represented in Figure 1. The viability threshold oxygen level in brain organoids *C_crit_*^17^ is also indicated in the graph. For a given radius, the minimum oxygen concentration in the gel-free constructs (blue) is higher than for the hydrogel-encapsulated ones (red). The radii at which the minimum oxygen concentration is equal to *C_crit_* determines the size limit for these systems, and corresponds to 712 μm for the gel-free models and 475 μm for the organoids surrounded by gel. On the other hand, comparing the green and red curves, it is clear that the use of a microfluidic channel can improve oxygenation within the constructs even in the presence of an ECM-mimicking shell. In fact, the maximum construct radius in the presence of a microchannel is 616 μm.

**Figure 2.**
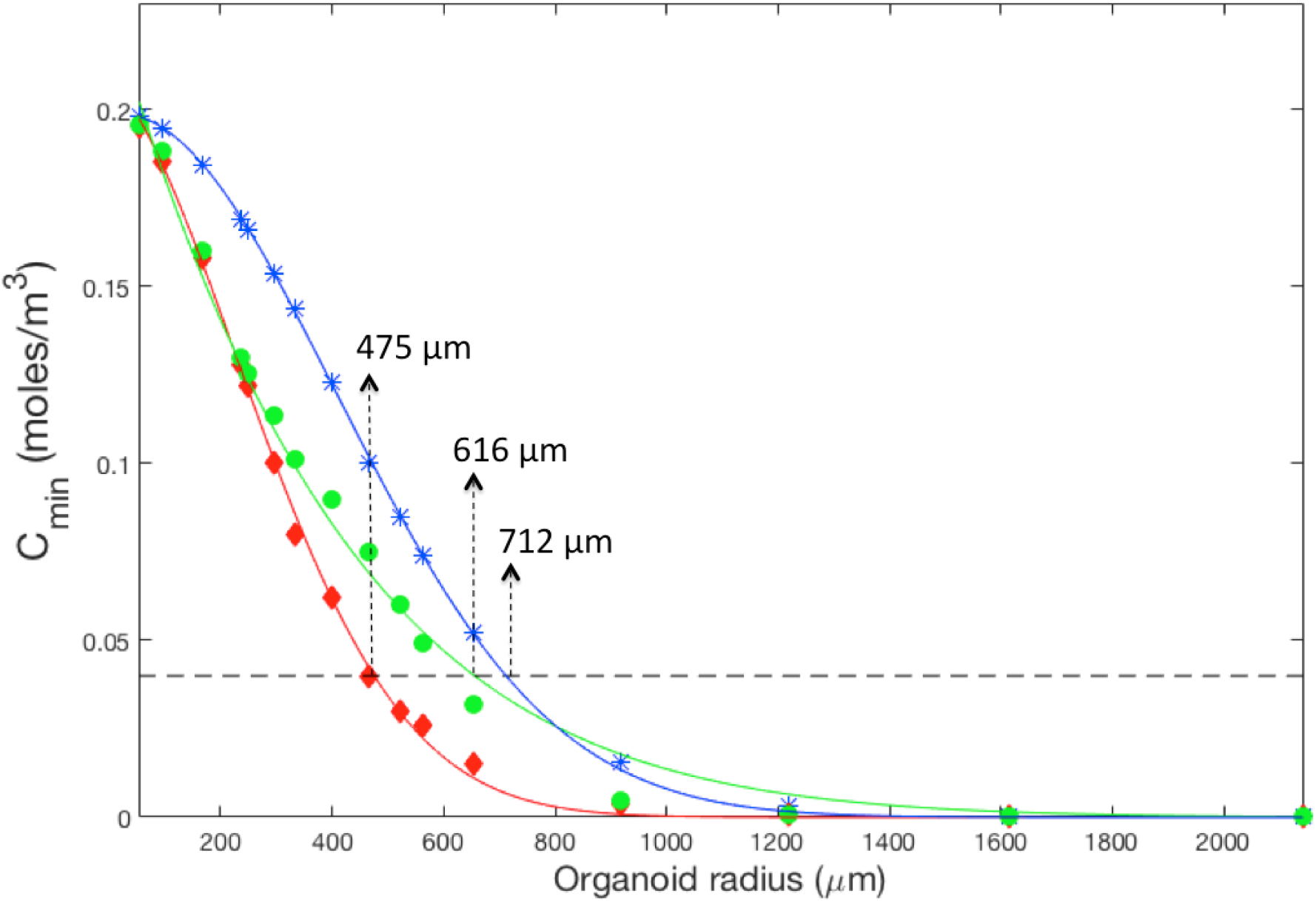
Minimum O_2_ concentration in constructs (*C_min_*) as a function of radius and corresponding fitted curves. Blue asterisks: gel-free organoids; red diamonds: organoids with gel shell; green circles: organoids with a gel shell and a microchannel. Fitting results: blue, R^2^:0.99; red, R^2^: 0.99; green, R^2^:0.98. The dotted black line represents *C_crit_*.

Table 4 summarizes the range of radii for which both criteria, b≈−1/4 and *C_min_*≥*C_crit_* are satisfied for each configuration presented in Figure 1. From the table it is clear that the working window for physiological relevance is very narrow for the organoid models sheathed by an ECM gel. Here, only constructs with radii between 236 and 475 μm satisfy both criteria. For the gel-free models, the range is even narrower, going from 654 to 712 μm. A much larger range of radii can be covered if a single microfluidic channel is incorporated in the organoids, as both b≈−1/4 and *C_min_*≥*C_crit_* are valid for the range 236 to 616 μm.

**Table 4.** Range of construct radii in which both KL law and oxygen concentration above the 0.04 mol/m^3^ threshold are maintained.

| | $r (c \geq C_{min})$ | $r(b \approx -1/4)$ | $r (C_{min} \geq C_{crit}) \cap r (b \approx -1/4)$ |
| --- | --- | --- | --- |
| **Gel-free, Figure 1A** | 54-712 | 654-2142 | 654-712 |
| **Gel shell, Figure 1B** | 54-475 | 236-562 | 236-475 |
| **Gel shell and channel, Figure 1C** | 54-616 | 236-2143 | 236-616 |

## Discussion

Developing physiologically relevant *in vitro* models to mimic cell and tissue organ response is one of the main challenges in experimental cell biology and tissue engineering. Organoid technology has emerged as a novel method to generate structural and functional tissue models^15,32–35^. Luminal organoids were first generated from intestinal stem cells by Clevers and coworkers^36^. As they are hollow, they are unlikely to suffer from necrotic cores^37^. On the other hand, non-luminal organoids such as brain and liver often possess non-viable centres, principally due to the lack of oxygen which is widely considered as the limiting metabolic resource for living cells^17,22^. Local oxygen levels also influence cell fate^38–42^, likely compromising tissue function and structural arrangement. Thus, further strategies to guarantee the nutritional needs of such organoids are required.

It has been suggested that allometric behaviour could be considered as a benchmark of physiological relevance^9,25^ and a recent study shows that KL can be achieved in high density constructs^12^. The characteristic features of non-luminal organoids, that is high cell densities and their potential to therefore follow physiological metabolic scaling as well as to adopt the structural and functional features of mammalian organs on one hand, and their tendency to develop an oxygen deprived core on the other, appear to be in contrast. In this context, our aim was to evaluate the minimum oxygen concentration and the *CMR* of non-luminal organoids to enable the design of cell culture protocols in which both quarter power scaling holds and the oxygen level in the organoids is greater or equal than the necrotic threshold of 0.04 mol/m^3^ (*C_crit_*). As illustrated in Figure 3, the intersection of the two criteria is argued to represent the window of physiologically relevant organoid sizes for designing constructs with better translational value. We therefore established 3D models of cell-filled spheres in a variety of configurations covering over 5 orders of magnitude of mass and including the range of sizes reported in the literature. The cell density and oxygen consumption parameters were those reported in the literature for brain organoids and stem cells respectively, while the boundary oxygen concentration was set at 0.2 mol/m^3^, as in ref.^12^. To identify the range of masses in which KL holds, the slope of log-log graphs of *CMR* versus mass was estimated. The masses and corresponding radii for which all the cells in the 3D constructs receive an adequate supply of oxygen were also computed.

**Figure 3.**
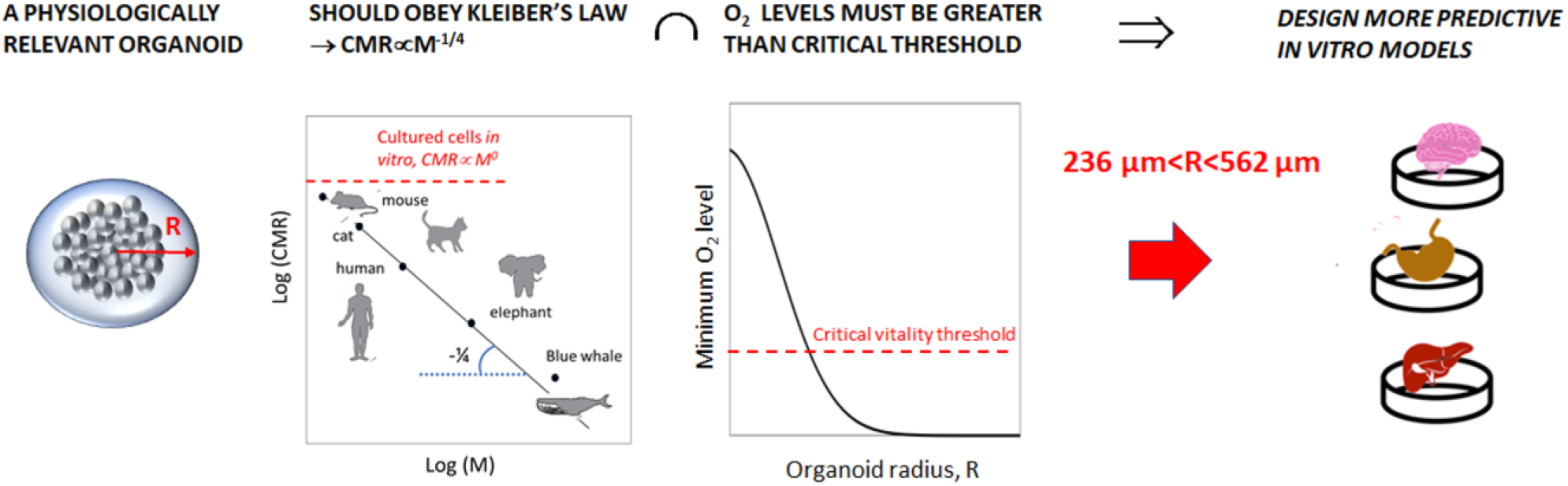
Physiologically relevant organoids should obey allometric scaling laws and cells in the centre of the constructs must be exposed to oxygen levels above the critical threshold for vitality. Using computational models to identify the range of radii which respect these two conditions, we can establish criteria for generating *in vitro* models with better translational value.

Some mini-organs, typically liver buds generated using mesenchymal cell-driven condensation, are dependent on an ECM-like substrate but do not require embedding in a gel to undergo differentiation and self-organisation^24^. Our results indicate that if cultured in the presence of a well-mixed and constant supply of freshly dissolved oxygen at the organoid-media boundary, these structures do not suffer from nutrient limitations over a wide range of masses (up to 1.5 mg, Table 4). The results are confirmed by experimental studies on liver buds by Ramachandran *et al*.^22^; the authors found no evidence of cell death in liver buds cultured in well-mixed fluidic bioreactors while those in static conditions developed a necrotic core. Mattei *et al*.^43^ demonstrated that the minimum oxygen level in liver buds with an average mass of a few mg cultured in bioreactors is 0.04 mol/m^3^, compared with 0.01 mol/m^3^ in static culture plates. As shown in Table 4, the gel-free structures resembling liver buds maintain the criteria of quarter power scaling (*b* ≈ −1/4) and viable oxygen concentrations only if their radii are maintained in a narrow range (between 654 and 712 μm in the models).

On the other hand, most of the current protocols for generating brain organoids rely on the use of ECM gels (e.g. Matrigel) which act as a 3D media supporting cell proliferation and differentiation. However, the gels also limit oxygen diffusion within the sample: indeed non-viable cores have been reported in these organoids. In fact, in the models the radii at which both the conditions (i.e. allometric scaling and minimum oxygen concentration above *C_crit_*) are satisfied are smaller than those for the gel-free constructs. Interestingly, in most studies, the sizes of organoids surrounded by a shell of ECM are generally smaller than those without a shell^13,14,22,35^.

We also developed a model to determine if the incorporation of a simple fluidic channel mimicking an elemental central blood vessel could widen the working window in which both criteria of *b*=−1/4 and *C_min_*≥*C_crit_* are satisfied (Figure 1C). The results show that the addition of the channel results in oxygen concentrations above *C_crit_*, and quarter power scaling for a wider range of radii, including those for brain organoids summarised in Table 1. Such channels have been reported in cortical organoids which are integrated into the host vasculature when implanted in animals^21^, but their fabrication *in vitro* remains a challenge.

It is important to highlight that we assumed a constant oxygen concentration at the outer surface of the geometries implemented, corresponding to the maximum dissolved oxygen concentration in water at 37°C at atmospheric pressure: this condition requires well-mixed and continuously renewed media through the use of fluidic devices such as bioreactors. Furthermore, although current technology using oxygen-sensitive fluorescent probes have made it possible to determine local intra-cellular oxygen levels^37,44^, accurate measurements of single cell and cell aggregate resource consumption rates are difficult to perform and scarce in the literature. The maximum oxygen consumption rate and the MM constant used here are based on published data for human mesenchymal stem cells. They may vary slightly from one cell population to another as different cells have different metabolic demands, and the critical range of radii for physiologically relevant organoid systems may vary accordingly.

Based on the considerations in this work and our previous study^12^, typical values of *b* and their significance in terms of resource availability are reported in Table 5. The exponent *b* is close to zero when all cells in the volume consume resources at the same rate because they all perceive the same oxygen concentration. As the constructs increase in size, cells on the periphery perceive and consume higher concentrations of oxygen than cells close to the centre, which adapt to lower levels of oxygen and hence lower consumptions rates - a characteristic of MM kinetics. As a consequence, the average *CMR* decreases and at the ‘knee’ of the log-log CMR versus mass graph, *b* is approximately −1/4 as in KL. At higher masses, as shown in ref.^12^, the coefficient *b* tends to −1/3, which indicates that resource uptake depends on the surface area of the construct^45^ and is diffusion limited.

**Table 5.** Typical values of the allometric exponents and their significance in terms of resource availability. (N.B. α=*b*+1).

| $\alpha$<br>($\text{BMR} \propto M^\alpha$) | $b$<br>($\text{CMR} \propto M^b$) | Notes |
| --- | --- | --- |
| 0 | -1 | Severe resource limitation |
| 2/3 | -1/3 | Surface area dependent consumption rate |
| 3/4 | -1/4 | Adaptive resource consumption rate (e.g. Michaelis Menten) |
| 1 | 0 | No resource limitation, maximal resource consumption rate |

In conclusion, we suggest that a resource-supplying network is not a necessary condition for quarter power scaling in high density organoids or cellular aggregates. However, despite obeying KL, non-luminal organoids may be resource limited as core oxygen levels fall below *C_crit_*. Thus, quarter power scaling in these structures is not a sufficient criterion for physiological relevance because part of the cell population may not perceive enough oxygen to guarantee their vitality. The range of sizes which guarantee physiological relevance is in fact quite limited and should be taken into consideration when employing organoids as mini *in vitro* models of human organs.

There is a great demand for improving organoid models with respect to their size, structure and composition: this study provides a first step towards the development of protocols for designing more predictive organoid systems based on objective criteria. Although current methods for integrating organoid and fluidic technology have focused on on-chip devices in which constructs are cultured inside fluidic channels^24^, we suggest that efforts be made to develop methods using bioprinting technology to fabricate organoids with a perfusable central^46^. The design of simple microcirculation-on-a-chip solutions must however be accompanied by accurate methods for regulating and monitoring oxygen and its consumption rates.

## Funding Statement

The work leading these results has received funding from the Fondazione Umberto Veronesi under the Post-Doctoral Fellowship 2018 and from the European Union’s Horizon 2020 research and innovation programme under Grant Agreement No 760813 (PATROLS). Funds from the Swiss National Science Foundation, through Project SINERGIA 2019 572 CRSII5_186422 /1 are also acknowledged.

## Authors Contributions

CM and AA: conception and design of the simulations. CM, AR and AA: interpretation of data and the drafting of the manuscript.

## Competing Interests

The authors have nothing to declare.

## Data Availibity

The equations and tables provided in the manuscript are sufficient to replicate the computations. Data files are available upon reasonable request.

## Notes

#### Summary of Updates

A new figure

