## Supplementary Materials for "Allometric Scaling of physiologically-relevant organoids"

A brief introduction to allometric scaling, the governing equations used in the finite element model and the mathematical methods used to determine the CMR are presented for the interested reader.

### Allometric scaling

Allometric scaling laws correlate the mass (*M* in kg) of an organism with any physiological parameter (*Y*) through an exponent ($\beta$) and a proportionality constant (*a*).

$$Y = aM^{\alpha} \qquad (1)$$

This exquisitely simple equation applies to all biological organisms because they are assembled according to the same basic design principles and using the same building blocks (mainly water) such that self-similarity is preserved across all scales [1]. The exponent $\beta$ is well known for a number of physiological parameters, such as heart rate ($\beta$=-1/4), basal metabolic rate or *BMR* ($\beta$=3/4) also known as Kleiber's Law, lifespan ($\beta$=1/4) [2]. While $\beta$ is considered a "universal" constant, the proportionality constant *a* is generally thought to hold for a given class, which in our case is Mammalia. The two constants, $\beta$ and *a* are obtained by plotting log-log graphs of the parameter *Y* against mass *M* for a few decades of mass. $\beta$ is the slope of the graph and *a* is the y-intercept.

For *BMR*, expressed as the moles of oxygen consumed per second, we can write Kleiber's Law:

$$BMR = aM^{3/4} \qquad (2)$$

On the basis that cell density (ρ, cells/$m^3$) is constant in all mammals [3], and that the density of biological organisms is close to that of water (Ω=1000 kg/$m^3$), the average metabolic rate per cell, or *CMR* in moles of oxygen/(cell.s) can be expressed as the total *BMR* divided by the number of cells (*#cells*) in the organism:

$$CMR = \frac{BMR}{\#cells} = \frac{a}{\rho}\Omega M^{-1/4} = a_{CMR} M^{-1/4} \qquad (3)$$

Here, $a_{CMR}$ is the new proportionality constant. The *CMR* versus mass relationship in cell cultures has been explored by West and co-workers, who deduced that in vitro cell cultures do not follow allometric scaling [4]. However, more recently Ahluwalia demonstrated that 3D cell cultures obey quarter power scaling if the oxygen concentration gradient within the 3D volume is sufficiently large [1].

**Governing equations**

For the simplest configuration (Figure 1S), the basic governing equation is the reaction-diffusion equation for oxygen, with Michaelis Menten type oxygen consumption. The convection term which considers a velocity vector $\underline{v}$ is also included for the sake of completeness.

$$\frac{\partial c}{\partial t} = D_{org}\nabla^2 c - \underline{v}.\nabla c - \frac{OCR \cdot \rho \cdot c}{k_m + c} \qquad (4)$$

Here, c is the oxygen concentration, $D_{org}$ is the diffusion constant within the organoid, $OCR$ is the maximum oxygen consumption rate per cell in the absence of resource limitation and $k_m$ is the Michaelis Menten constant.

Assuming steady state, zero velocity field and spherical symmetry, the equation reduces to

$$\frac{D_{org}}{r^2}\left(r^2\frac{dc}{dr}\right) = \frac{OCR \cdot \rho \cdot c}{k_m + c} \qquad (5)$$

Table 3 in the main text reports the values of the constants used. The boundary conditions are: a concentration of $c=c_o=0.2$ mol/m$^3$ at the surface of the sphere, which implies a continuous and well-mixed resource supply. Equation 5 cannot be solved analytically, but the concentration field within a spherical volume of radius R can be computed using finite element methods (we used COMSOL Multiphysics version 4.3 COMSOL AB, Stockholm, Sweden). $C_{min}$ and the radius at which $c=C_{crit}$ can thus be identified.

The surface integral function in COMSOL can be used to calculate the number of moles/s which cross the surface of the sphere. The inward flux ($-J = D\nabla c$ at $r=R$) represents the number of moles of oxygen crossing the surface and entering into the sphere per unit time and area. Its integral over the surface of the sphere is thus the total metabolic rate or BMR:

$$BMR = \iint_A D\nabla c \cdot dA \qquad (6)$$

Which for spherical symmetry can be expressed as:

$$BMR = D\left.\frac{dc}{dr}\right|_R 4\pi R^2 \qquad (7)$$

From equation 3, the CMR is simply the BMR divided by the number of cells in the sphere, is:

$$CMR = \frac{\iint_A D\nabla c \cdot dA}{\# cells} \qquad (8)$$

To derive the value of the allometric exponent $\beta$ for CMR, we plot a log-log graph of the computed CMR against mass ($M = \Omega \frac{4}{3}\pi R^3$) for spheres of different sizes and determine the slope as described in the main text.

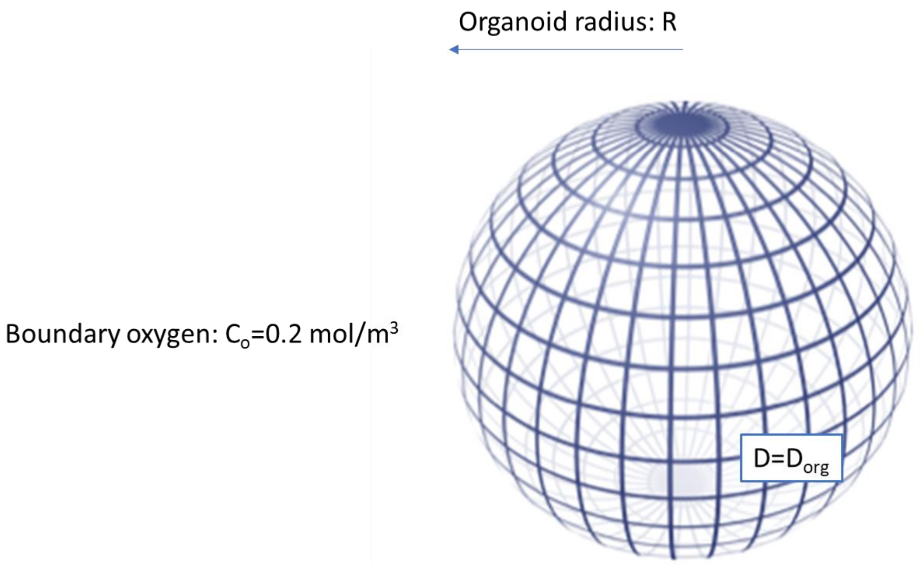


***Figure 1S: The simplest configuration: a sphere with radius R.***

The second configuration, illustrated in Figure 2S also considers a 3D medium with a diffusion constant $D_M$ surrounding the cells, which required for the generation of stem-cell derived organoids. Ignoring the convection term, considering steady state conditions and spherical symmetry, the equations are:

$$\frac{D_{org}}{r^2}\left(r^2\frac{dc}{dr}\right) = \frac{OCR \cdot \rho \cdot c}{k_m + c} \text{ , } 0<r<R$$

$$0 = \frac{D_M}{r^2}\frac{d}{dr}\left(r^2\frac{dc}{dr}\right) \text{ , } R<r<R+\delta$$

and continuity at the boundary r=R, while at r=R+δ( R), $c = c_o = 0.2\ mol/m^3$

The hydrogel volume is constant, therefore the thickness $\delta$ is a function of R.

The *CMR* and related allometric exponent are obtained as above.

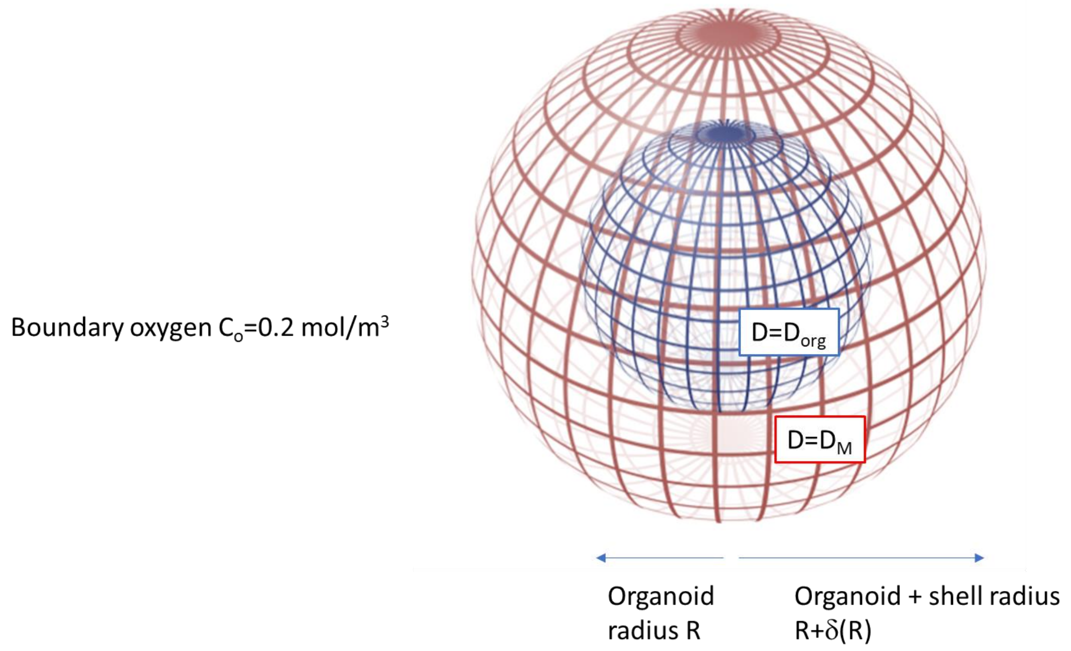


***Figure 2S: The second configuration: a sphere with radius R surrounded by a gel of thickness $\delta$***

Finally, the microfluidic configuration with an elementary vessel through the centre of the sphere through which water-based medium flows is represented in Figure 3S. This configuration has 3 domains with the following equations.

$$0 = D_{org}\nabla^2 c - \frac{OCR \cdot \rho \cdot c}{k_m + c} \qquad \text{in the blue domain}$$

$$0 = D_M \nabla^2 c \qquad \text{in the red shell domain}$$

$$\left.\begin{aligned} 0 &= D_{water}\nabla^2 c - \underline{v}.\nabla c \\ \Omega \frac{D\underline{v}}{Dt} &= -\nabla P + \mu \nabla^2 \underline{v} \\ 0 &= \nabla.\underline{v} \end{aligned}\right\} \qquad \text{in the green cylinder domain}$$

The conditions pertaining to the external boundaries are illustrated in the figure, while continuity is maintained at all inner boundaries. When calculating the *CMR*, the surface integrals of the inward flux at the surface of the inner (oxygen consuming) and outward flux from the surface of the central cylinder are added together.

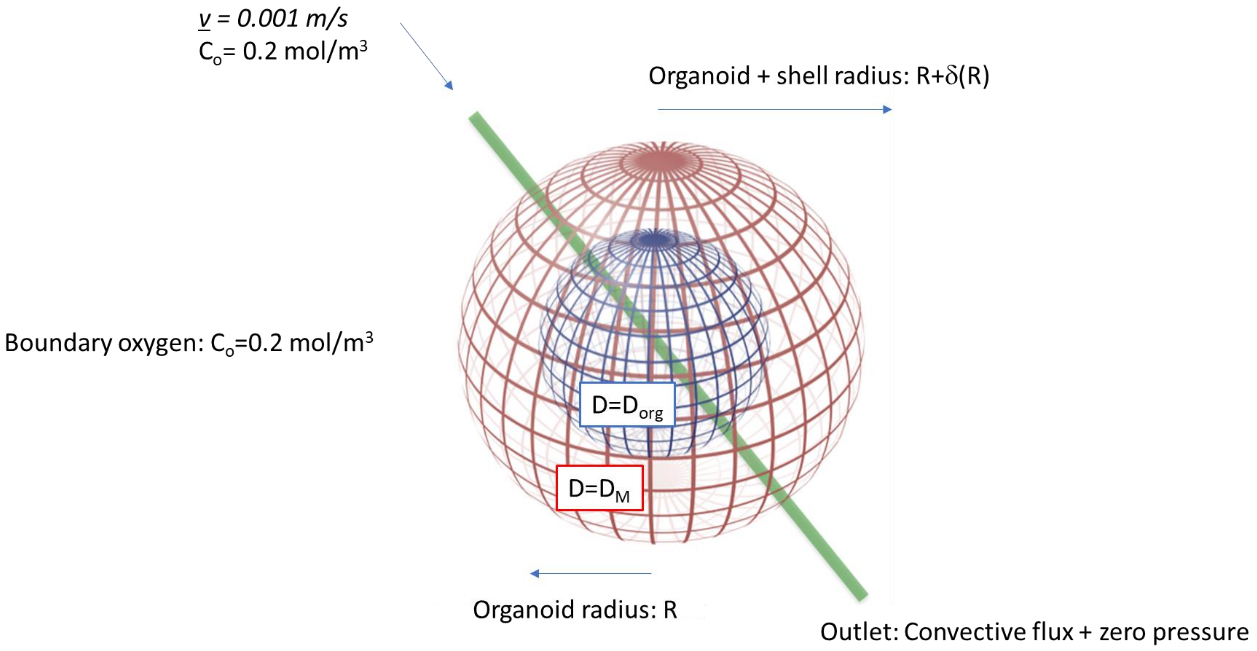


***Figure 3S: The third configuration: a sphere with radius R surrounded by a gel of thickness $\delta$ and with a central a microfluidic channel (length 4 cm, radius 10 μm).***
